# Unveiling a Novel Memory Center in Humans: Neurochemical Identification of the *Nucleus Incertus*, a Key Pontine Locus Implicated in Stress and Neuropathology

**DOI:** 10.1101/2023.09.08.556922

**Authors:** Camila de Ávila, Anna Gugula, Aleksandra Trenk, Anthony J. Intorcia, Crystal Suazo, Jennifer Nolz, Julie Plamondon, Divyanshi Khatri, Lauren Tallant, Alexandre Caron, Anna Błasiak, Geidy E. Serrano, Thomas G. Beach, Andrew L. Gundlach, Diego F. Mastroeni

## Abstract

**Background:** The *nucleus incertus* (NI) was originally described by Streeter in 1903, as a midline region in the floor of the fourth ventricle (4V) of the human brain with an ‘unknown’ function. More than a century later, the neuroanatomy of the NI including its forebrain target regions has been described in lower vertebrates, but not in humans. Therefore, we examined the neurochemical anatomy of the human NI using several markers, including the neuropeptide, relaxin-3 (RLN3), and began to explore the distribution of the NI-related RLN3 innervation of the hippocampus.

**Methods:** Histochemical staining of serial, coronal sections (30 µm) of control human postmortem pons was conducted to reveal the presence of the NI by detection of immunoreactivity (IR) for the neuronal marker, microtubule-associated protein-2 (MAP2), two markers present in rat NI, glutamic acid dehydrogenase (GAD)-65/67 and corticotrophin releasing hormone receptor 1 (CRHR1), and RLN3, which is highly expressed in a major population of NI neurons in diverse species. *RLN3* and vesicular GABA transporter 1 (*vGAT1*) mRNA was detected by multiplex, fluorescence in situ hybridization. Postmortem pons sections containing the NI from an Alzheimer’s disease (AD) case were immunostained for phosphorylated-tau (AT8 antibody), to explore potential relevance to neurodegenerative diseases. Lastly, sections of human hippocampus were stained to detect RLN3-IR and somatostatin (SST)-IR, as SST is expressed in interneurons targeted by RLN3 projections in rodents.

**Results:** In the dorsal, anterior-medial region of the human pons, neurons containing RLN3– and MAP2-IR, and *RLN3/vGAT1* mRNA-positive neurons were observed in an anatomical pattern consistent with that of the NI in other species. GAD65/67– and CRHR1-immunopositive neurons were also detected within this area. Furthermore, RLN3– and AT8-IR were co-localized within NI neurons of an AD subject. Lastly, RLN3-IR was detected in neurons within the CA1, CA2, CA3, and DG areas of the hippocampus, in the absence of *RLN3* mRNA. In the DG, RLN3– and SST-IR were co-localized in a small population of neurons.

**Conclusions:** Aspects of the anatomy of the human NI are shared across species, including a population of RLN3-expressing neurons and a RLN3 innervation of the hippocampus. Accumulation of phosphorylated-tau in the NI suggests its possible involvement in AD pathology. Further characterization of the neurochemistry of the human NI will increase our understanding of its functional role in health and disease.

Graphical Abstract *Created with BioRender.com*

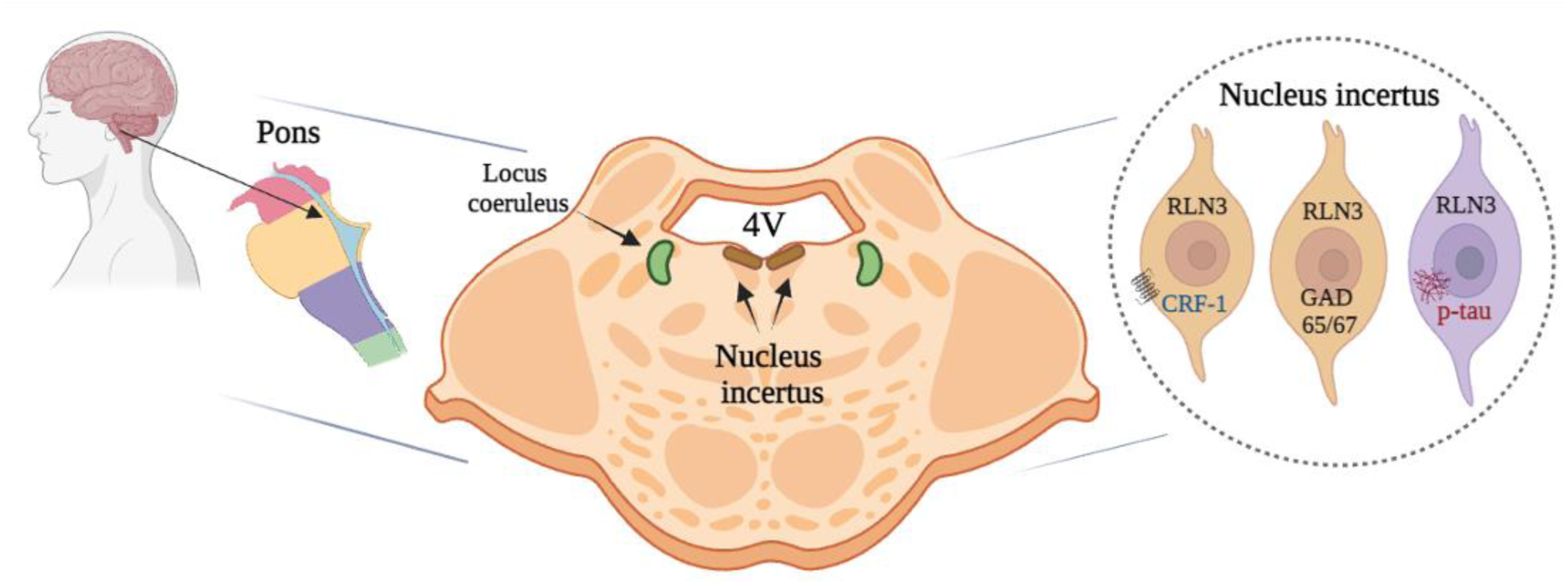

## Background

The brainstem receives a range of sensory and autonomic inputs from the periphery and the spinal cord, and neurons in this region project throughout the brain to distribute these integrated signals to influence multiple higher-processing networks. Ascending networks including the locus coeruleus (LC) and medullary raphe nuclei are highly susceptible to neurodegenerative diseases, including AD [1–4]. In addition, extensive studies of the LC have contributed to understanding the involvement of the hippocampus in memory [5–7].

The brainstem NI or the ‘uncertain nucleus’, was originally described by anatomist, George L. Streeter in 1903 as a midline region in the floor of the 4V of the human brain with ‘unknown’ function [8]. An equivalent area was also described in guinea pig (1926) [9], in hamster (1954) [10], and in cat (1968) [11]. Thereafter, many decades later, a large majority of neurons in the rat NI or *nucleus O* [12] was shown to express the inhibitory neurotransmitter, γ-aminobutyric acid (GABA), as reflected by staining against the GABA-synthesizing enzyme, GAD, and in situ hybridization (ISH) for *vGAT1* mRNA [13, 14]. Since then, the neuroanatomy of the NI and its connections throughout the brain has been systematically mapped using various histological methods in rats [14–19], mice [20–24], zebrafish [25], and in non-human primate [26], but not in humans.

In general, the NI is a bilateral nucleus, located in the midline adjacent to the more lateral LC [15, 17, 22, 24]. Its anatomy varies slightly in different species. In adult rats, the NI extends for ∼0.7 mm from Bregma −9.12 mm to Bregma −9.84 mm and is referred as ‘nucleus O’ by Paxinos and Watson [12, 17, 19]. In mice, the NI is located more ventral to the 4V than in rats, in the midline central gray of the dorsal pons [22]. In the macaque, the NI is located adjacent to the 4V, within the ventromedial central gray of the pons/medulla, and medial to the LC [26].

Recent research has identified specific neuroanatomical and functional connections between the NI and the septohippocampal system (SHS) and demonstrated that the NI plays a role in special and contextual fear memory [14, 21, 23, 27, 28]. The predominant inhibitory neurotransmitter system (i.e., GABAergic system) is associated with bi-directional projections between the NI and SHS [29, 30], and these NI-related circuits can regulate fear memory via effects on SST-positive interneurons in the hippocampus and hippocampally-projecting septal neurons [21, 23]. In addition, the degeneration of GABA and SST/GABA neurons occurs in the medial SHS and basal forebrain cortical systems in AD and other neurogenerative dementia [31, 32].

Importantly, a major population of NI neurons has been demonstrated to produce RLN3 in multiple species [24, 26, 33, 34]. RLN3 is a highly-conserved neuropeptide [35, 36] that signals via the relaxin-family peptide-3 receptor (RXFP3)[24, 37, 38]. RLN3/RXFP3 signaling is strategically positioned to modulate SHS-related learning and memory processes that are integral in AD symptomology [30, 39, 40]. In addition, NI is a stress responsive region and its neurons express receptors for the stress hormone, corticotropin releasing hormone (CRH) [41–43]. Thus, in light of the likely importance of the NI and associated GABAergic and neuropeptide systems in memory and key sensory and autonomic processes in humans under different physiological conditions, it is timely to investigate the neurochemical anatomy of human NI neurons.

## Methods

### Postmortem human tissue

The postmortem human brain tissue used in this study was fully characterized and generously provided by the Arizona Alzheimer Disease Research Center (ADRC) Pathology Core, Banner Sun Health Research Institute, Brain and Body Donation Program (BBDP, Sun City, AZ, USA, http://www.brainandbodydonationprogram.org). Autopsy and tissue fixation were conducted as described [44]. Details for each subject are provided in Table 1.

**Table 1.**
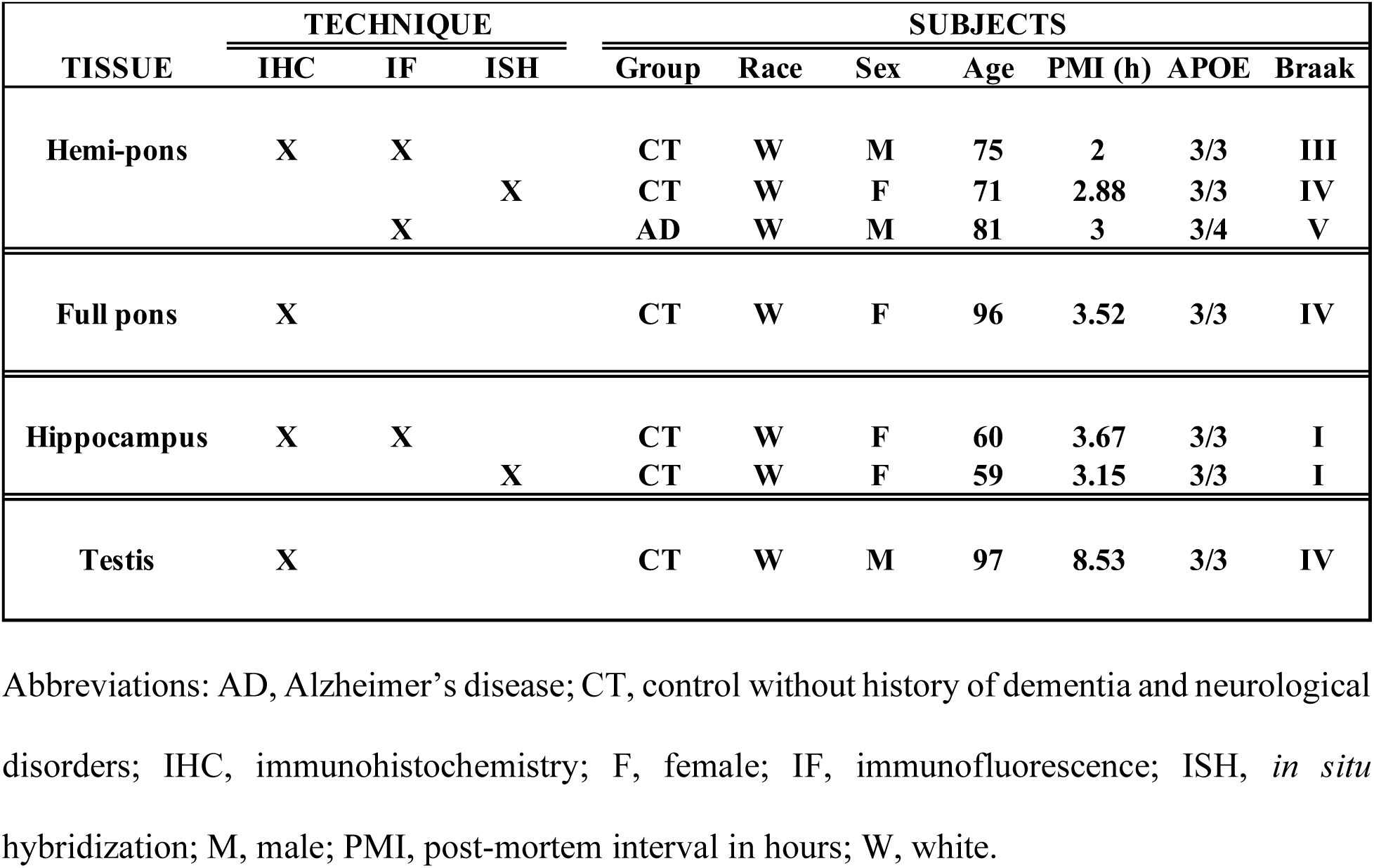
Clinical and pathological characteristics of studied cases for each anatomical region and technique used.

### Multiplex fluorescence in situ hybridization

Expression of *RLN3* and *vGAT1* mRNA in postmortem human pons and hippocampus (from control subjects; see Table 1) was assessed using RNAscope™ multiplex fluorescence ISH (Advanced Cell Diagnostics (ACD), Hayward, CA, USA). Briefly, fresh-frozen, coronal sections (10 μm) were cut at –20°C using a cryostat, and mounted onto Superfrost-Plus slides, according to the BBDP protocol [44]. All procedures were conducted following the user manual for RNAscope™ Multiplex Fluorescent V2 assay (fresh-frozen sections) provided by the manufacturer (ACD). The slides were stored at –80°C until a 1 h fixation in a freshly-prepared solution of 4% formaldehyde in phosphate-buffered saline (PBS, pH 7.4, initially 4°C) at room temperature (RT), followed by washing in PBS and dehydration in ethanol solutions of increasing concentration (50, 70, and 100%). Dehydrated sections were stored at –20°C over weekend and then air-dried, outlined with Immedge Hydrophobic Barrier Pen (Vector Laboratories, Burlingame, California, USA) and incubated with hydrogen peroxide for 10 min at RT and washed in distilled water. Next, after applying protease IV pre-treatment solution (ACD) for 30 min at RT and washing in PBS, the sections were hybridized for 2 h at 40°C with a solution of Multiplex probes for: *RLN3* (Hs-RLN3-C1, Cat. No. 590151, ACD) and *vGAT1* (Hs-SLC32A1-C3, Cat. No. 415681-C3, ACD). Sections were washed in 1×Wash Buffer (ACD) between every hybridization step. Following all amplification steps, HRP-C1 (labelled with TSA Vivid Fluorophore 650, 1:1500 in TSA buffer) and HRP-C3 (labelled with TSA Vivid Fluorophore 570, 1:1500 in TSA buffer) signal was developed. Finally, the tissue was counterstained with DAPI, coverslipped with ProLong Gold antifade reagent (Invitrogen, Thermo Fisher Scientific, Life Technologies Corporation, Eugene, OR, USA) and imaged using an Axio Imager M2 fluorescent microscope (Zeiss) with an automatic stage and Axiocam 503 mono camera (Zeiss). 20×/0.5 EC Plan Neo-Fluar objective was used for acquisition of panoramic z-stack images of the NI and the hippocampus areas (scaling: 0.227 μm in x and y, and 1.250 μm in z axis) and 40×/1.3 Oil EC Plan Neofluar objective for obtaining single representative z-stack images of NI neurons (scaling: 0.114 μm in x and y, and 0.280 μm in z).

The images were then processed in Zen software (3.3 blue edition and 2.3 SP1 black edition, Zeiss) and ImageJ [45] to improve the signal-to-noise ratio and converted into maximum intensity projection images.

### Immunohistochemistry (IHC)

Chromogenic immunohistochemical studies were completed using sections from human pons, hippocampus, and testis (see Table 1). Tissue was fixed according to the BBDP protocol [44], sectioned at 30-40 µm on a microtome, and mounted on charged glass slides for histology. Briefly, free-floating sections were blocked in 1% H_2_O_2_ and 3% bovine serum albumin (BSA, Cat. No. D5637, Sigma-Aldrich, St. Louis, Missouri, USA). Pons sections were stained with Hematoxylin and Eosin (H&E, Cat. No. 26041-06, EMS, Hatfield, PA, USA and Cat. No. E511-100, Thermo Fisher Scientific, Tempe, AZ, USA). Sequential sections were incubated in antisera raised against MAP2 (1:8,000, Cat. No. ab183830, anti-rabbit, Abcam, Fremont, CA, USA) or RLN3 (1:1,500, Cat. No. PA47448, anti-goat, Invitrogen, Carlsbad, CA, USA), overnight (ON) at 4°C. Sections were washed and then incubated in species-specific secondary antisera (1:1,000 – Cat. No. Vector BA-1000, goat anti-rabbit IgG (H+L), Biotinylated, Vector Laboratories Inc., Burlingame, California, USA, or Cat. No. Vector BA-5000, rabbit anti-goat IgG (H+L), Biotinylated, Vector Laboratories Inc., Newark, California, USA) for 2 h at RT. Sections were washed and incubated in 1:1,000 avidin/biotin reagent (VECTASTAIN ABC-HRP Kit (Standard), Cat. No. PK-4000, Vector Laboratories Inc., Burlingame, California, USA), washed and incubated in Tris buffer 0.05M (20 mM Tris-HCl and 150 mM NaCl, Sigma-Aldrich, Cat. Nos. T5941, T1503, S3014, St. Louis, Missouri, USA), 3,3′-Diaminobenzidine (DAB, Cat. No. D5637, Sigma-Aldrich, St. Louis, Missouri, USA; Concentrated: 10 mg/mL), saturated nickel ammonium sulfate solution (Alfa Aesar, Cat. No. A18441, Ward Hill, MA, USA), and 1% H_2_O_2_ (Starting reagent: Hydrogen peroxide solution 30%, Cat. No. 216763, Sigma-Aldrich, St. Louis, MO, USA). Sections were reacted for 10 min (MAP2) and 30 min (RLN3), dried, dehydrated through graded alcohols, cleared in xylene, and mounted using Permount (Cat. No. 17986-01, EMS, Hatfield, PA USA). Hippocampus and testis sections were only incubated with RLN3 antibody and subjected to the same steps described above.

#### Image analysis

Imaging of tissue sections stained using immunohistochemistry was performed using an Olympus IX70 microscope equipped with bright field illumination (Olympus, Tokyo, Japan). Results were captured using an Olympus DP-71 color digital camera. Whole slide images were acquired using an Olympus VS200 slide scanner and processed using Olympus DESKTOP v3.4.1 to generate representative images.

### Immunofluorescence

Postmortem brain samples were selected to best match critical factors such as postmortem interval (PMI), age, and sex, as well as other relevant covariates (Table 1). Sections were washed (3×) in phosphate-buffered saline with 0.1% Tween^®^ 20 detergent (PBS-T, Cat. No. 20012-027, GIBCO PBS pH 7.2 (1X), Life Technologies Corporation, Grand Island, New York, USA, with Tween^®^ 20 detergent, Cat. No. P1379, Sigma Aldrich Company, St. Louis, Missouri, USA), and blocked with 3% BSA, with incubation for 1 h. After further washing, sections were incubated in a range of combined primary antisera [RLN3 (1:1,500, Cat. No. PA47448, anti-goat, Invitrogen, Carlsbad, CA, USA) and AT8 (1:500, MN1020, anti-mouse, Invitrogen, Carlsbad, CA, USA), or CRHR1 (1:500, Cat. No. ab150561, anti-rabbit, Abcam, Fremont, CA, USA), or GAD65/67 (1:500, Cat. No. PA5-104543, anti-rabbit, Invitrogen, Carlsbad, CA, USA), or MAP2 (1:500, Cat. No. ab183830, anti-rabbit, Abcam, Fremont, CA, USA), or SST (1:200, Cat. No. ab108456, anti-rabbit, Abcam, Fremont, CA, USA)] antisera; ON at 4°C.

Sections were washed 3× in PBS-T and incubated in species-specific, fluorophore-conjugated secondary antibodies (1:1,000, Red: Cat. No. A11036, AlexaFluor 568 goat anti-rabbit IgG (H+L) 2 mg/mL, and 1:1,000, Green: Cat. No. A11034, AlexaFluor 488 goat anti-rabbit IgG (H+L) 2 mg/mL, or Cat. No. A11055, donkey anti-goat IgG (H+L) 2 mg/mL, or Cat. No. A11029, goat anti-mouse IgG (H+L) 2 mg/mL, Life Technologies, Eugene, OR, USA). After a final wash, sections were mounted, dipped in 1% Sudan Black (Cat. No. 199664, Sigma-Aldrich, St. Louis, Missouri, USA to reduce autofluorescence, and coverslipped with UltraCruz containing DAPI (UltraCruz® Hard-set Mounting Medium, sc-359850, Dallas, TX, USA).

#### Imaging analysis

Imaging of tissue sections stained with fluorescent markers was performed using an Olympus IX70 microscope equipped with epifluorescence illumination. Images were captured using an Olympus DP-71 color digital camera. High-resolution micrographs were acquired using a Leica SP8 confocal microscope (Leica, Wetzlar, Germany).

### Specificity of RLN3 antiserum – Dot blot and pre-adsorption tests

Dot blot analysis was used to test the specificity of the polyclonal RLN3 antibody used in the immunohistochemical studies for recognizing human RLN3 peptide. Lyophilized human RLN3 (Cat. No. 035-36, Phoenix Pharmaceuticals, Burlingame, CA, USA) was reconstituted in distilled water and was used at a concentration of 1 µg/µL. Two µl of RLN3 peptide were added to the center of a 0.2 μm nitrocellulose membrane (Cat. No.1620146, Bio-Rad, Hercules, CA, USA). Non-specific sites were blocked by incubation with 5% BSA in Tris-buffered saline with 0.1% Tween^®^ 20 detergent (TBS-T, 20 mM Tris-HCl and 150 mM NaCl, Sigma-Aldrich, Cat. Nos. T5941, T1503, S3014, with Tween^®^ 20 detergent, Sigma Aldrich, Cat. No. P1379, St. Louis, Missouri, USA) for 1 h at RT, and then the membrane was incubated with RLN3 antiserum (1:15,000, Cat. No. PA47448, anti-goat, Invitrogen, Carlsbad, CA, USA) for 30 min. The membrane was washed 3 × 10 min in TBS-T, incubated with a secondary antibody conjugated with HRP (1:500, Cat. No. PI-9500, Horse Anti-Goat IgG (H+L), peroxidase; Vector Laboratories Inc., Newark, CA, USA) for 30 min at RT, then washed 3 × 10 min in TBS-T and 1 × 5 min in TBS. The membrane was incubated with ECL reagent for 1 min, then imaged on an Amersham Imager 680 (GE), with an exposure time of 90 s.

Validation of the Dot blot result was also performed using a pre-adsorption assay. An aliquot of RLN3 antiserum (see details in IHC section) was pre-incubated ON with RLN3 peptide (molar ratio 1:10 antibody:peptide). RLN3 immunohistochemistry was repeated as described above in 40 µm, free-floating sections from postmortem human testis (a positive control tissue for RLN3 peptide expression [46].

## Results

In this study, we identified the NI of the human brain, in postmortem tissue using RLN3, MAP2, GAD 65/67, and CRHR1 as neurochemical markers for the area. Combining techniques that preserve the neuroanatomy – ISH and IHC – we first detected the presence of *RLN3* mRNA and peptide in the human NI, using hemi-brain sections. The neuronal distribution within the dorsal pons at the level of the NI was first mapped using an antibody for MAP2, which is expressed in neurons [47] (**Fig. 1A, B**). Multiple neuronal populations were stained for MAP2, in the dorsal, anterior-medial region of the pons and in the area adjacent to the 4V and dorsal to the medial longitudinal fasciculus (mlf); and the distribution was similar to the distribution of neurons found in the area of macaque brain identified as the NI [26]. Once a neuronal population at the anticipated rostrocaudal level of the human NI was identified, RLN3-IR was detected to confirm the location and better define the anatomical boundaries of the NI (**Fig. 1C**). The presence of *RLN3* mRNA was detected to investigate whether RLN3 translation was also occurring within the NI (**Fig. 2A**). *RLN3* mRNA and peptide were confirmed within the anteromedial level of the pons. Importantly, as in other species tested [37], ISH results indicated that *RLN3* mRNA is colocalized with *vGAT1* mRNA in human NI (**Fig. 2 B, C**), which indicates the GABAergic nature of NI neurons synthesizing RLN3 in the human brain. In line with findings in rats [17, 20], and to some extent in mice [20, 24], the anatomy of the NI in humans was characterized by a compact and more dispersed regions. Previous reports in the rat brain [17] indicated that the *pars compacta* displayed a dense population of neurons near the 4V, while the *pars dissipata* contained fewer, more sparse neurons along the superior part of the mlf.

**Fig. 1.**
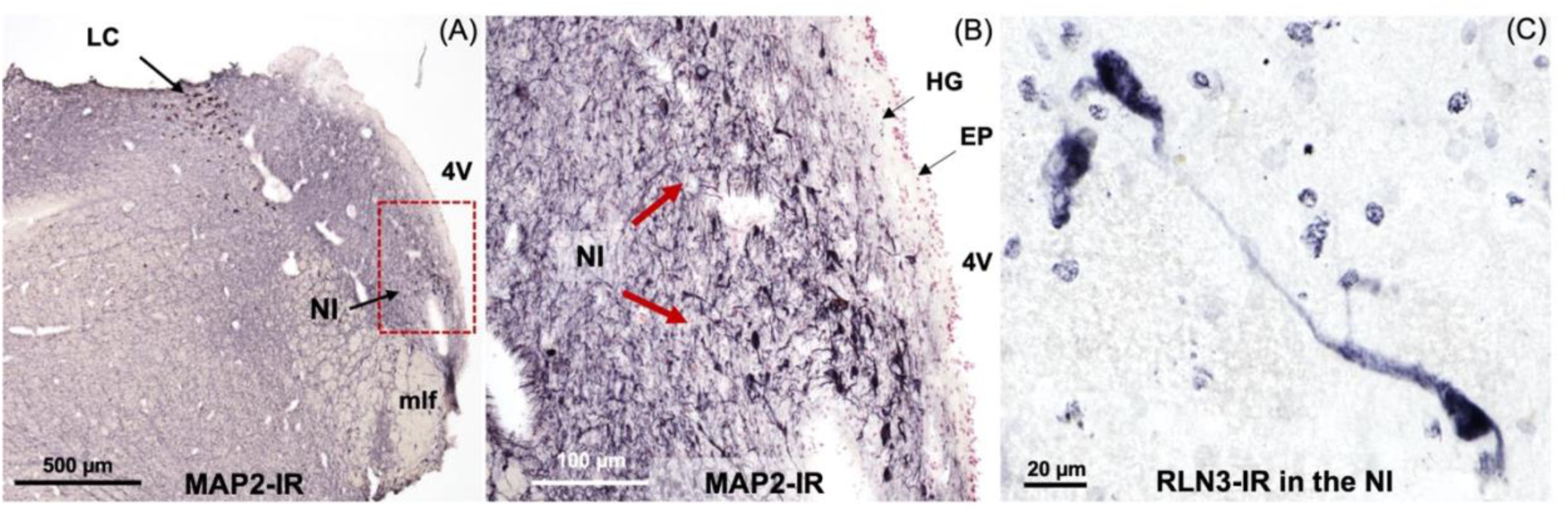
RLN3-Immunoreactivity in the Coronal Anterior Hemi-Pons of Human Brain. (**A**) Neuronal populations in the human pons, stained for MAP2-IR. MAP2 is a marker of neuronal cells, their perikarya and dendrites [88]. (**B**) A higher magnification view of the human pons, with red arrows indicating MAP2-IR neurons in an area adjacent to the medial longitudinal fasciculus equivalent to that containing RLN3-positive NI neurons in the macaque. (**C**) RLN3-immunopositive neurons in human NI. Abbreviations: EP, ependyma, 4V, fourth ventricle, HG, Hypocellular gap, IR, immunoreactivity, LC, locus coeruleus, MAP2, microtubule-associated protein 2, mlf, medial longitudinal fasciculus, NI, nucleus incertus.

**Fig. 2.**
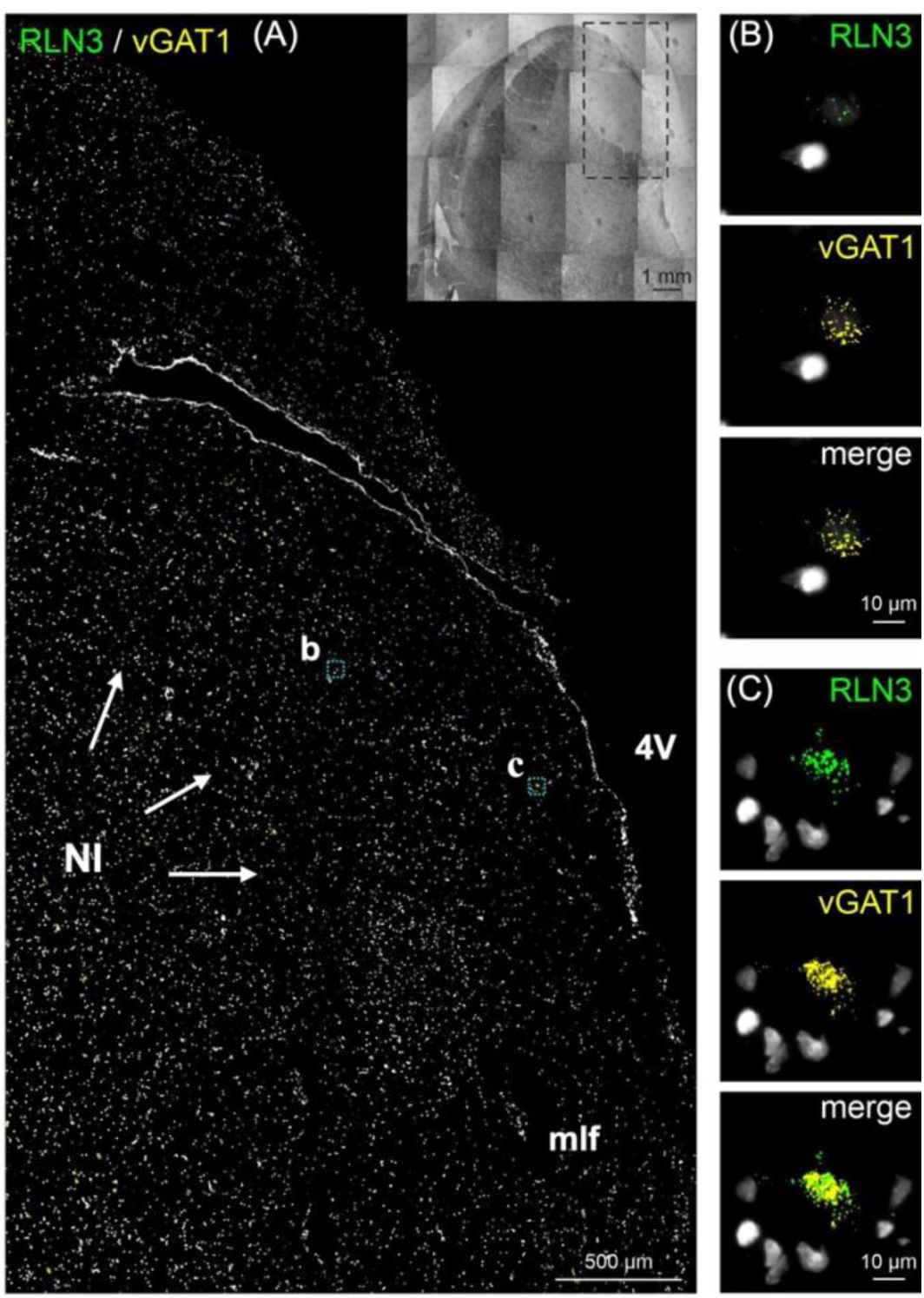
*In Situ* Hybridization (RNAscope™) Detection of *RLN3* mRNA in the Coronal Anterior Hemi-Pons of the Human Brain. (**A**). **(B)** Higher magnification images of *RLN3* mRNA (green particles) and (**C)** *vGAT1* mRNA (yellow particles) co-localized in neurons in the NI area (merge). Inset represents the hemi-pons section under brightfield illumination. Abbreviations: 4V, fourth ventricle, mlf, medial longitudinal fasciculus, NI, nucleus incertus, *vGAT1*, vesicular GABA transporter-1.

We validated the specificity of the RLN3 antisera for detection of human RLN3 using sections of postmortem human testis (positive control)[46] incubated with or without RLN3 antisera subjected to pre-adsorption of the native peptide (**Supplementary** Fig. 1A-F). The pre-adsorption assay completely abolished specific immunostaining. In addition, the dot blot experiment demonstrated that the RLN3 antisera binds to native RLN3 peptide (**Supplementary** Fig. 1G), although we did not assess its cross reactivity with other related or unrelated peptides. In studies to confirm the neuronal nature of RLN3 immunopositive cells, we co-incubated RLN3 and MAP2 antisera (**Fig. 3**) and immunofluorescent detection revealed consistent colocalization of these markers within neurons in the NI region, while in areas outside the NI, MAP2 positive neurons were negative for RLN3-IR (**Fig. 3**).

**Fig. 3.**
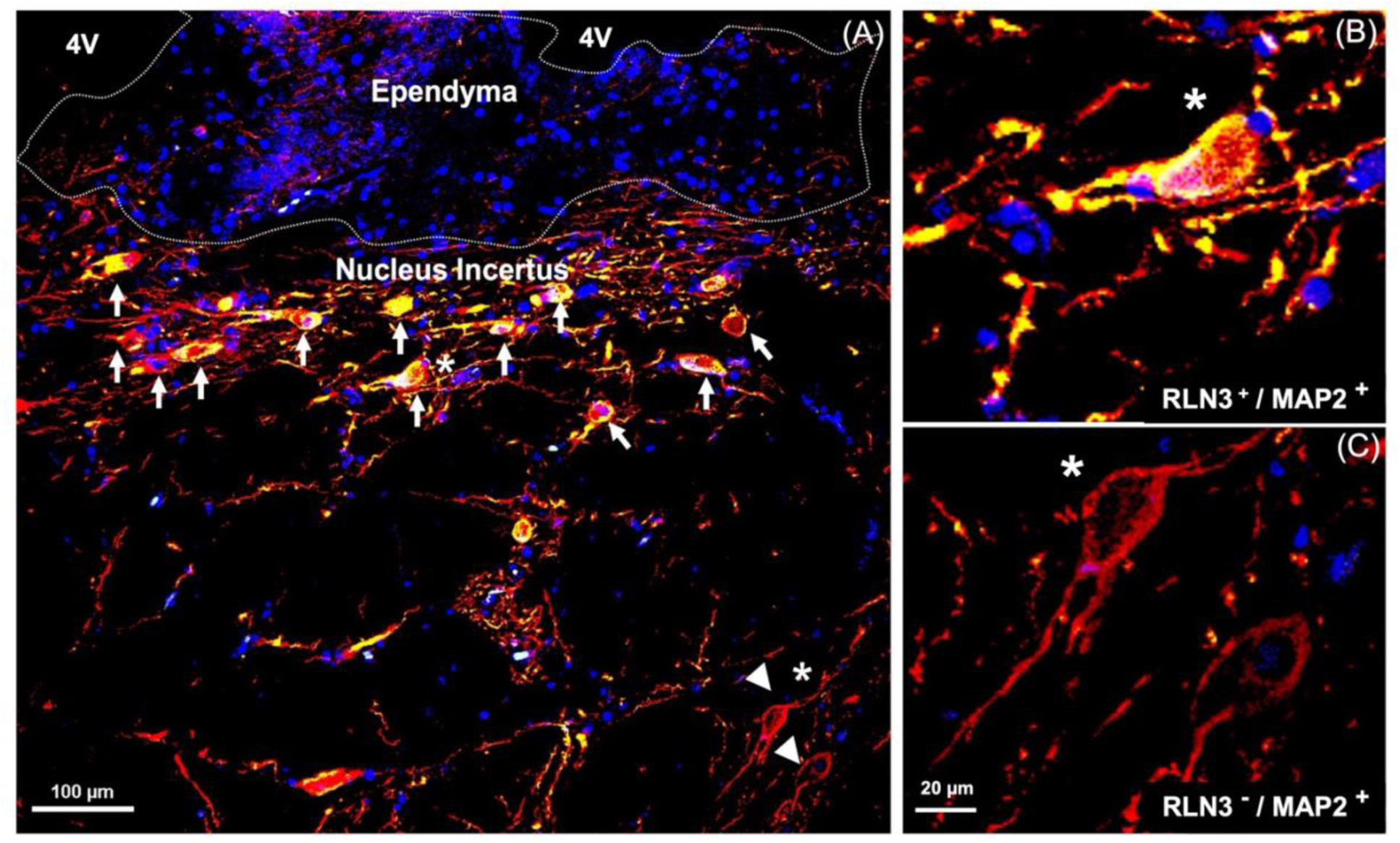
Relaxin-3 is expressed in neurons. Microtubule-associated protein 2 is a marker of neuronal cells, their perikarya, and dendrites [88]. (**A)** NI in humans; white arrows indicate RLN3-immunoreactivity co-localizing with MAP2. (**B)** Higher magnification view of a RLN3^+^/MAP2^+^ neuron indicated with a white star. (**C)** Higher magnifications of RLN3^-^/MAP2^+^ neurons indicated in A the white head arrows. MAP2 (red), RLN3 merging with MAP2 (yellow), DAPI (blue). Abbreviations: 4V, fourth ventricle, MAP2, microtubule-associated protein 2, RLN3, relaxin-3.

After identifying the human NI region in sections of hemi-brain samples and validating the ability of the RLN3 antiserum to recognize the native peptide, we conducted further experiments in sections of intact, bilateral pons and immunostained them for additional markers. We confirmed the nuclear and cytoplasmic structures using H&E (**Supplementary** Fig. 2); the distribution of MAP2-containing neurons in the same NI area identified using hemi-pons sections (**Fig. 4**); the presence of the NI bilaterally was revealed by detection of RLN3-IR (**Fig. 5**); and different (light/heavy) staining of the peptide was observed in different neurons within the area (**Fig. 5D-E**).

**Fig. 4.**
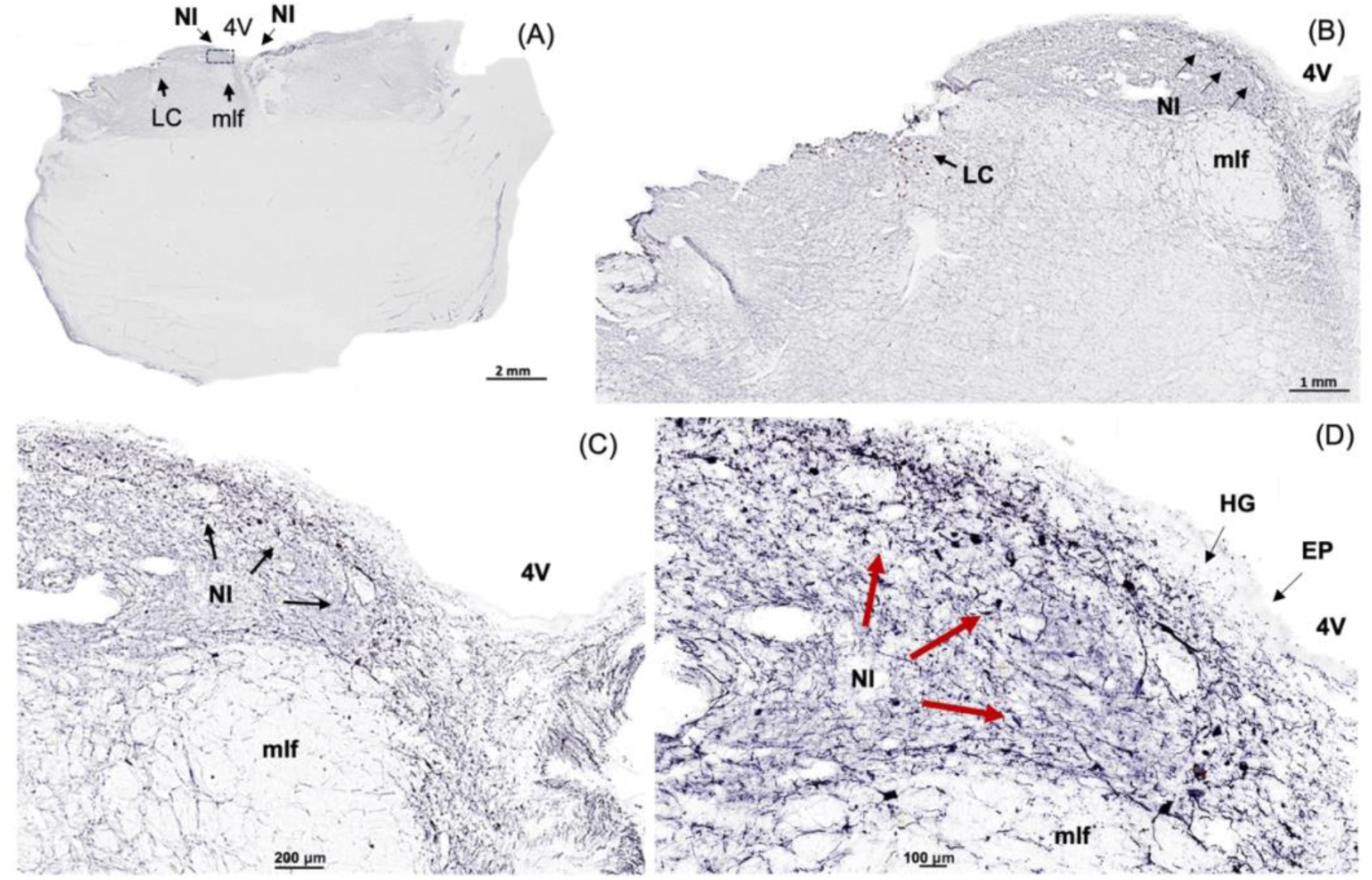
Microtubule-associated protein 2 Immunoreactivity in Neurons in the Coronal Anterior Pons of Human Brain. (**A)** Low-power image of the neuroanatomy at the human pons containing the NI stained for MAP2-IR, a marker of neuronal perikarya and dendrites [88]. (**B)** Higher magnification image of the boxed area in A. (**C)** Higher magnification of the boxed area in B. (**D)** Higher magnification of the boxed area in C. Abbreviations: EP, ependyma, HG, Hypocellular gap, IR, immunoreactivity, LC, locus coeruleus, MAP2, microtubule-associated protein 2, mlf, medial longitudinal fasciculus, NI, nucleus incertus, 4V, fourth ventricle.

**Fig. 5.**
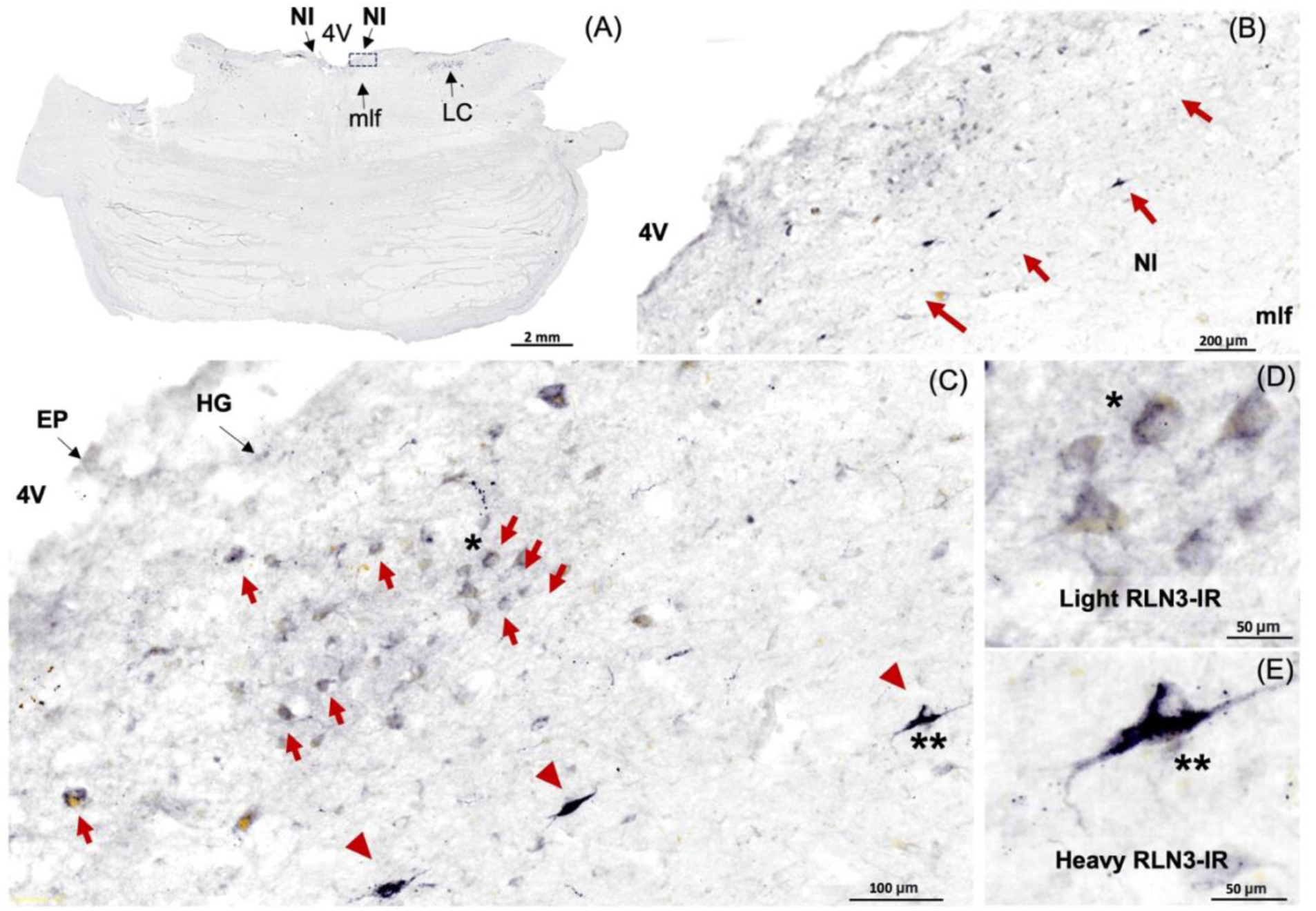
RLN3-Immunoreactivity in the Coronal Human NI. (**A)** Low power image of the anterior level of the pons, with the NI located at the floor of the 4V, above the mlf and lateral to the LC. (**B)** Higher magnification images of the boxed area in A. (**C)** Higher magnification images of the boxed area in B, with neurons lightly and heavily stained for RLN3 indicated by red arrows and red arrowheads, respectively. (**D, E)** Higher magnification images of neurons with light and heavy staining for RLN3-IR, respectively. Abbreviations: EP, ependyma, 4V, fourth ventricle, HG, Hypocellular gap, IR, immunoreactivity, LC, locus coeruleus, mlf, medial longitudinal fasciculus, NI, nucleus incertus.

Additional immunostaining was completed to further verify possible similarities in the neurochemistry of the human NI with other species. In all non-human species examined (i.e., rats, mice, macaque), RLN3-positive neurons are GABAergic [14, 33] and express the CRHR1 in the rat [34, 48]. GAD65/67– and CRHR1-IR were detected in the human NI (**Fig. 6**), and both CRHR11– and GAD65/67-IR were co-localized with RLN3-IR (**Fig. 6C, F**).

**Fig. 6.**
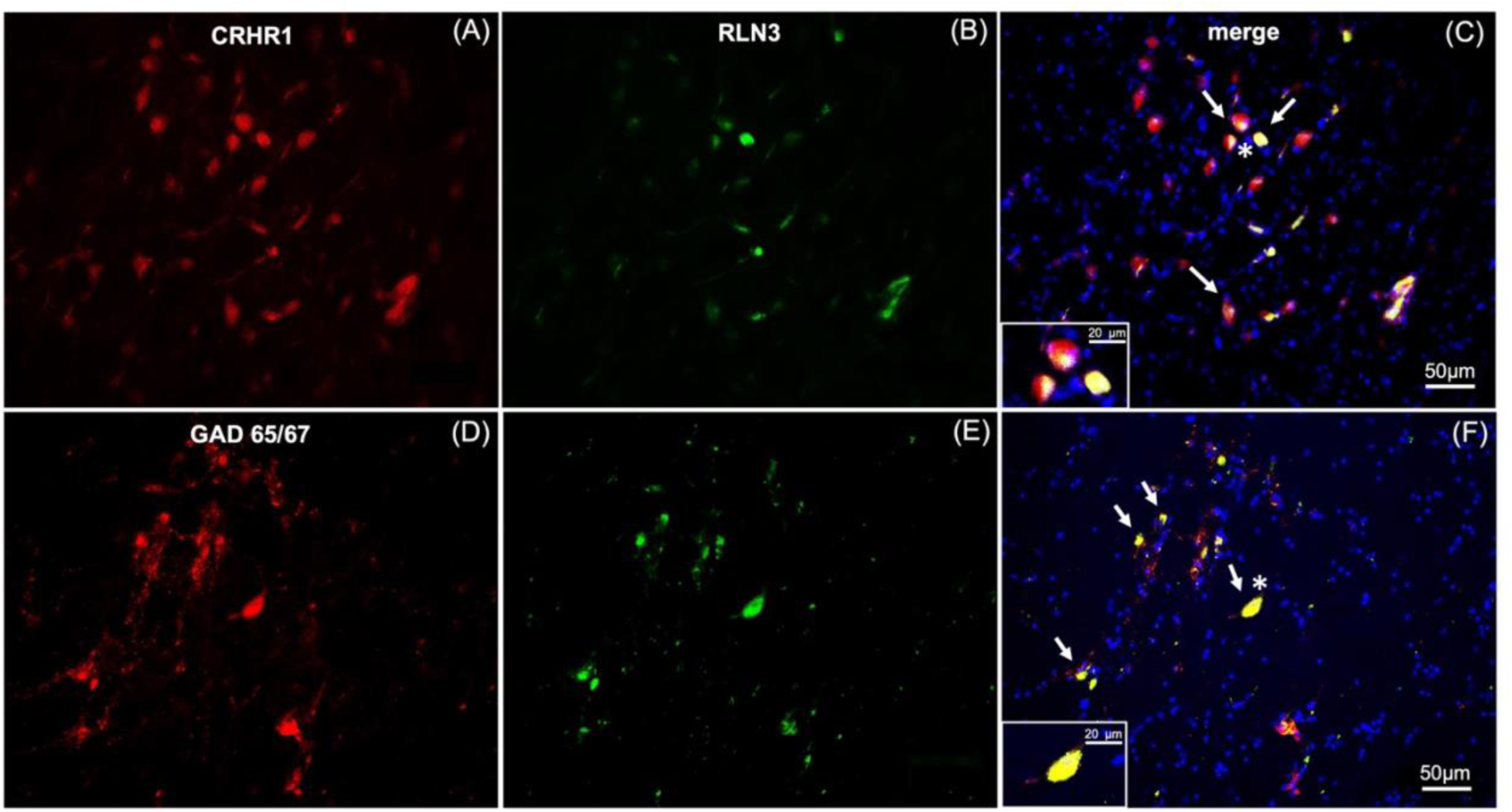
RLN3-Immunoreactive Neurons in the Coronal Human NI Co-Express CRHR1– and GAD 65/67-immunoreactivity. (**A-C)** RLN3 positive neurons and surrounding cells in the NI region, co-express corticotropin releasing hormone receptor 1 immunoreactivity (CRHR1; white arrows). (**D-F**) RLN3 positive neurons co-express glutamic acid decarboxylase 65/67 immunoreactivity (GAD65/67, white arrows). Insets represent higher magnification images of neurons s indicated by an asterisk in C and F. CRHR1 and GAD 65/67 (red), RLN3 (green), DAPI (blue), merged image (yellow).

In light of the possible association between the NI neural network and memory and cognition under normal and pathological conditions [21, 23, 39, 40], we examined the relationship between a marker for AD-related pathology, phosphorylated-tau and RLN3-expressing neurons in the NI from an AD subject (**Fig. 7**). Notably, phosphorylated-tau was detected at the NI level of pons sections from an AD subject, as reflected by AT8-IR and these neurons also contained RLN3-IR (**Fig. 7C**), whereas no AT8-IR was detected in NI neurons of a control subject (**Fig. 7A**). Successful detection of AT8-IR in LC of AD sections was used as a positive control for the antibody (data not shown).

**Fig. 7.**
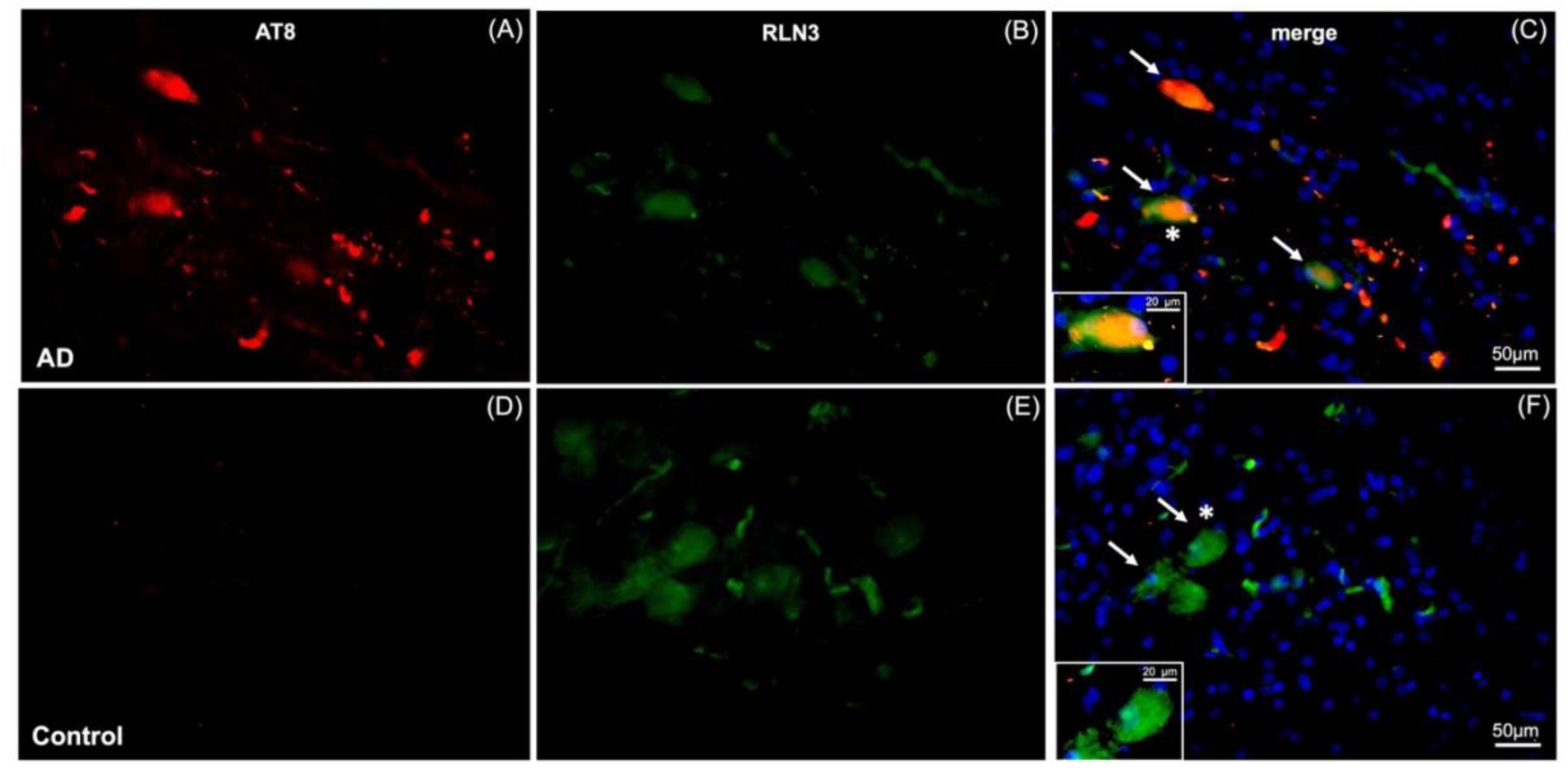
Neurofibrillary Tangles Co-localized with RLN3-Immunoreactivity in the Coronal NI of Human AD Brain. (**A-C)** AT8 (a marker for neurofibrillary tangles, NFT) and RLN3^+^ staining of neurons in the NI of an AD case, with co-expression NFT (indicated by white arrows). (**D-F)** AT8 and RLN3 staining of the NI of a control, age and sex-matched, non-demented subject. Insets in C and F represent higher magnification images of neurons marked with asterisk, and reveal the co-localization of NFT and RLN3 in AD but not control brain. AT8 (red), RLN3 (green), AT8 and RLN3 (yellow), DAPI (blue).

Lastly, we conducted initial studies of the presence of NI-related RLN3-IR, in postmortem sections of hippocampus from a control subject, and observed a distinct accumulation of RLN3-IR in sparse neurons in the CA1-2 and CA3/DG layers (**Fig. 8**). Notably, RLN3 mRNA was not detected within the hippocampus (data not shown), indicating peptide accumulation in the area.

**Fig. 8.**
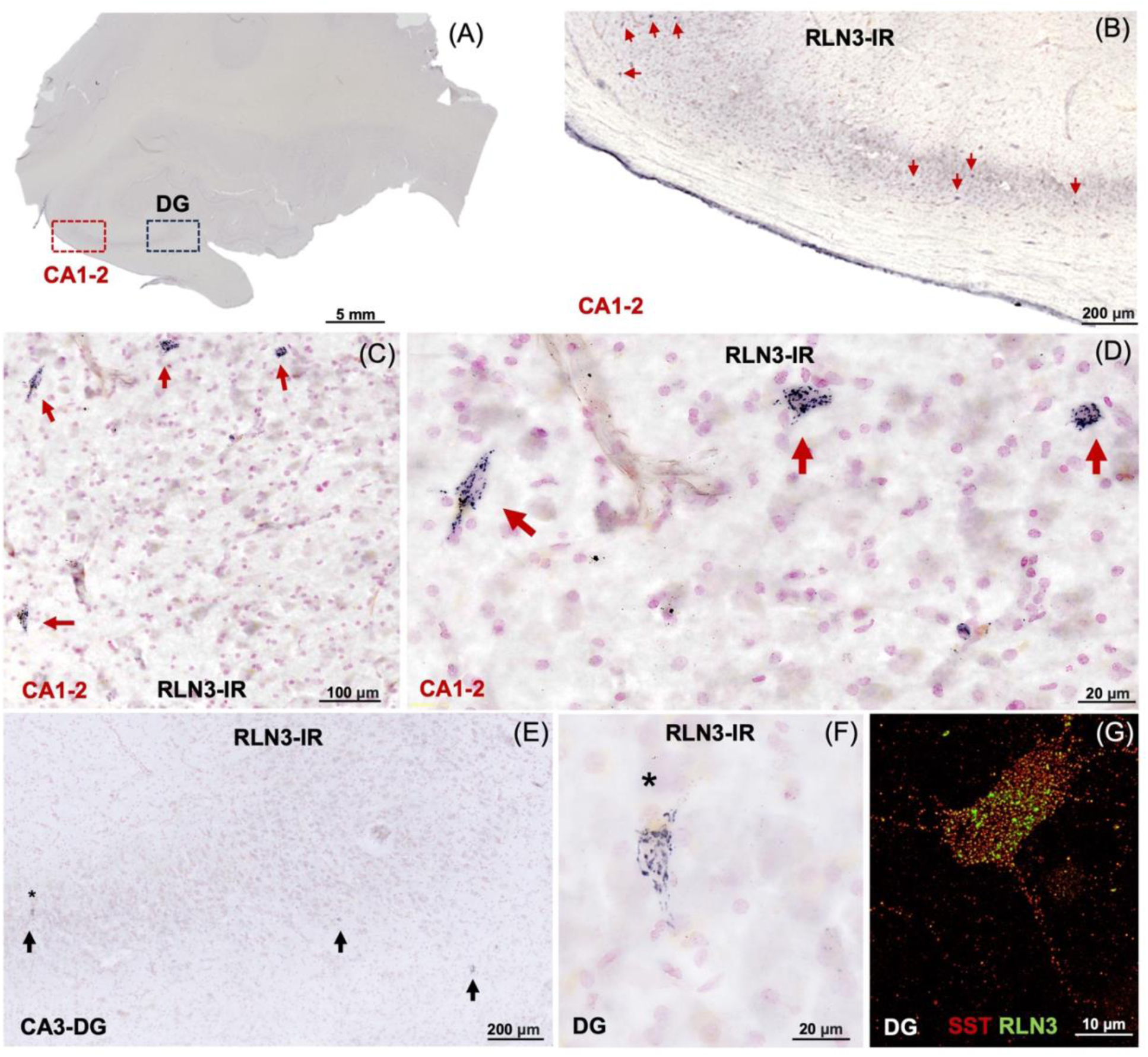
RLN3-Immunoreactivity (IR) Within Human Hippocampal Neurons. (**A**) Overview of the neuroanatomy of the coronal hippocampus lightly counterstained with neutral red. (**B-D)** Higher magnification images of RLN3-immunopositive cells within the CA1-2 (red boxed area in A). (**E, F)** High magnifications of RLN3-IR in the cells within the CA3 and dentate gyrus (DG, black boxed area in A). (**G)** Interneuron within the DG labeled for RLN3-IR (green) and somatostatin-IR (SST, red). The pattern of staining suggests an accumulation of the neuropeptide on the surface of these neurons, which express RLN3 receptor, RXFP3 in mice [76].

## Discussion

The results of the current study indicate that aspects of the anatomy and the neurochemistry of the NI are preserved across species and that the human NI shares similarities with non-human primates such as the macaque [26] and common experimental species, such as rats and mice [19, 37, 49]. Marker proteins of the GABAergic and CRH-mediated stress response systems, and phosphorylated-tau were detected within human RLN3 NI neurons in control and AD subjects, respectively.

Numerous earlier anatomical studies have characterized the connectivity and neurophysiological activity of the NI in rat and mouse brain, including the strong projections to the SHS system, and to multiple cortical and limbic areas involved in emotional cognition [14, 17, 18, 21], while others have characterized the nature of the transmitters, neuropeptides, and receptors expressed by different populations of NI neurons in these species [14–22], which provide an insight to the likely functional role of the NI in the human brain.

In the following sections, the significance of the current results is reviewed in the context of previous preclinical studies. It includes an overview of the existing knowledge on the role of the NI, and in particular RLN3 signaling systems, in spatial and contextual fear memory, and stress. Finally, it addresses the limitations of the current study and future experiments that are warranted based on our important initial findings.

### Identification of the NI in human brain

In the current study, we used MAP2-IR to detect neuronal populations in an area in the human dorsomedial pontine tegmentum equivalent to that reported to contain the NI in vertebrates [17, 22, 26]. After using MAP2 to identify a neuronal population appropriately positioned adjacent to the floor of the 4V and medial to the LC, we used serial, free-floating formalin-fixed hemi-brain sections to detect the presence of RLN3, a neuropeptide marker for the NI.

### Neurochemical profile of the NI and its functional implications

In rats and mice, RLN3 is primarily expressed in the NI [24, 34–36, 38] with some additional positive cells being found in the pontine raphe nucleus (PRn), the lateroventral periaqueductal gray (PAG), and a region dorsal to the substantia nigra (SN) pars compacta [33, 34]. We observed high densities of RLN3-IR in the human NI, similar to that observed in other species [26, 50]. We also detected neurons expressing low density of RLN3-IR. Further investigation is needed to understand whether RLN3-IR neurons in the hippocampus might accumulate the peptide via receptor-mediated and might show correlation with the low RLN3-IR density observed in the NI.

RLN3 is co-expressed with GAD– and GABA-IR in GABAergic projection neurons in rats [14, 33, 51]. In humans, we also observed GAD65/67-IR co-localized with RLN3-IR and mRNA encoding the vGAT1 co-localized with *RLN3* mRNA. In rats and mice, populations of NI neurons express other peptides, including neuromedin-B (NMB) [20] and cholecystokinin (CCK) [14, 18] and RLN3 is co-localized with NMB in the mouse [14, 22], but in a separate population to the CCK neurons in rat [14, 22]. However, unlike RLN3, which is confined to small populations of neurons in the pontine raphe, PAG, and dorsolateral to SN [24, 33, 34], NMB and CCK are more widely expressed throughout neurons in the forebrain [52, 53]. Furthermore, multiple anatomical and pharmacological studies in the rat have revealed modulatory inputs to the NI from a range of other key neuropeptide systems, including CRH, orexin/hypocretin [54, 55] and melanin-concentrating hormone (MCH) [56], as reflected by expression of CRHR1 [48] orexin/(hypocretin) receptors (OX2, OX1) [54, 55, 57], and MCH receptor-1 (MCH-1) [56], further reflecting the importance of the NI in the control of arousal and attention [58], processes highly related to cognition. NI neurons in rats and mice also express receptors for the monoamine transmitters, serotonin [serotonin-1A receptor (5-HT_1A_)] [59] and dopamine [D2-like receptor (D2R)] [14, 60]. Several of these putative influences have been explored experimentally and found to affect the activity of rat NI neurons [19, 43, 55, 56, 60, 61]. For example, pre-clinical studies using behavioral challenges, pharmacological interventions, lesions, and electric manipulations in rats, revealed that the NI was activated by stress and is a component of the stress response/initiation circuits [24, 34, 42, 48, 62]. In the current study, CRHR1-IR was detected within human NI neurons, aligned with these pre-clinical findings [19, 34], and stress is known to affect memory at multiple points of the neuraxis [63]. In addition, anterograde tracing in mice revealed that CRH neurons in the dorsomedial medulla innervate the NI and have been shown to regulate key aspects of sleep that drive memory consolidation [64].

In this regard, the SHS is a major target of studies aimed at understanding the etiology of age– and dementia-related memory decline; and studies in animal models have revealed that the RLN3/NI network shares anatomical similarities with other neuromodulatory networks implicated in the control of memory and arousal, and AD symptoms, such as the serotonergic, cholinergic, and noradrenergic systems [21, 65–70]. Indeed, extensive studies of the LC have contributed to understanding the involvement of the hippocampus in memory [71]. Importantly, stress and CRH also affect the activity of these systems [72–74]

Notably, the NI is the source of an ascending GABAergic pathway to the SHS, and NI neurons display activity related to hippocampal theta rhythm in rats and mice [15, 19–21, 48, 64, 75]. Moreover, it has recently been shown that the NI plays a role in fear memory formation in mice via direct inhibition of the SST neurons of the hippocampus [21, 23]. This evidence confirmed earlier proposals, based on several studies in rodents, that demonstrated the involvement of the NI-RLN3/RXFP3 system in declarative and contextual memory [19, 27, 28, 75]. These recent studies employing optogenetics and viral-based pathway mapping revealed that the ascending arm of the NI-SHS pathway can regulate fear memory via effects on SST-positive interneurons in the hippocampus and on hippocampally-projecting septal neurons [21, 23], and that NI activity can be regulated by descending inputs from the prefrontal and retrosplenial cortex [23]. In this regard, earlier studies have reported that SST interneurons receive a RLN3 innervation and express RLN3 receptors (RXFP3 mRNA) [51, 76, 77].

Notably, the degeneration of GABA and somatostatin/GABA neurons occurs in the hippocampus and basal forebrain-cortical systems in AD and other neurodegenerative dementias [78]. Therefore, overactivity or degeneration of the inputs from the NI may have adverse consequences for memory formation and may be a target for neural dysregulation by AD pathological processes. In our initial studies, we observed an accumulation of RLN3-IR in a distinct population of neurons in the CA1, CA2, and DG layers of the human hippocampus. In the DG, RLN3-IR was present in neurons containing SST-IR, and this peptide ‘accumulation’ may be associated with the normal function or pathological dysfunction of the RLN3/RXFP3 system at different stages of aging or age-related pathology.

### Possible impact of AD pathology on NI RLN3 neurons

Brainstem networks including those ascending from the LC and the raphe nuclei are highly susceptible to neurodegenerative diseases, including AD [79–82], and phosphorylated-tau has been used as a marker of neurodegeneration in diseases such as Alzheimer’s Disease and related Dementias (ADRD) [83]. Interestingly, phosphorylated-tau deposition was demonstrated at the level of the pons in cases diagnosed with AD [84]. In fact, pioneering studies by Braak and colleagues report that intracellular pretangles are first identified in the LC and various other brainstem nuclei, years before the presence of mature tangles in the limbic system. Notably, we observed an accumulation of phosphorylated-tau (reflected by AT8-IR) in RLN3-containing neurons of the NI of a Braak stage V AD subject, suggesting a possible association of the NI with AD etiology.

A likely role of NI and related circuits in memory formation [21, 40], and evidence that tau accumulation induces synaptic and spatial memory impairment [85, 86], highlights the importance of further characterizing the neurochemical anatomy of the human NI, and the impact of AD on this profile.

#### Limitations of these studies

While postmortem human brain tissue has provided valuable insights into the molecular profile of human NI neurons, it is important to acknowledge that this approach is not able to capture dynamic changes that occur in the brain of living individuals. In addition, sampling from hemi-brains may introduce some degree of bias, as it does not sample the complete bilateral NI. While efforts have been made to ensure accurate and representative sampling, future studies could consider adjusting the analysis to account for potential differences between sampled hemi-brains and the complete bilateral NI.

Postmortem tissue from elderly patients presents neuronal loss. Postmortem tissue from young adults would be beneficial to quantify the neuronal population in the NI. While free-floating formalin-fixed sections present a well conserved neuroanatomy and low levels of autofluorescence, availability of this specific type of tissue is reduced. Comparative studies using larger sample sizes would be warranted. Finally, we acknowledge that all the subjects used in this study were Caucasian, which may limit the generalizability of our findings to other racial and ethnic backgrounds. It is important to recognize the potential influence of genetic and environmental factors that can vary across different populations. To address this limitation, future studies should include a more diverse cohort of subjects to ensure the broad applicability of results. Institutions that serve more diverse populations could assist the recruitment of participants from various racial and ethnic backgrounds.

#### Future experiments

Further characterization of the molecular profile of neurons within the human NI as a function of age and AD is now warranted, and additional studies should consider employing techniques that allow real-time monitoring of human NI activity, such as functional imaging methods [87]. Such approaches would provide a more comprehensive understanding of the temporal dynamics of human NI function as well as the cellular underpinnings.

## Conclusions

The NI is an important but understudied nucleus in the anterior pons of the human brain. RLN3 peptide is the fingerprint of the NI. Accumulation of phosphorylated-tau was found in the NI of an AD patient. RLN3 peptide was detected in a population of hippocampal neurons, in the absence of its corresponding mRNA. In order to understand the involvement of the NI and related neural circuits in memory and ADRDs, future comparative studies using imaging and postmortem tissue should be conducted.

## Declarations

### Ethics approval and consent to participate

Written informed consent for autopsy was obtained in compliance with institutional guidelines of Banner Sun Health Research Institute. The Institutional Review Boards for Banner Sun Health Research Institute approved the operations of the Brain and Body Donation Program, including recruitment, enrollment, autopsy and sharing of biospecimens and data. In addition, samples were analyzed anonymously (e.g. sample numbers) throughout the experimental process.

## Consent for publication

Not applicable.

## Availability of data and materials

Not Applicable.

## Authors’ contributions

C. A. and A.L.G. conceived the study and designed the experiments. C.A., A.G., A.T., A.I., J.P., C.S., J.N., L.T., D.K., performed the experiments. C.A., D.F.M, G.E.S., T.G.B., A.B., A.C. and A.L.G. analyzed the data. D.F.M., T.G.B., G.E.S, A.B. and A.L.G. supervised the research. C.A. wrote the manuscript and prepared the figures, with editorial input from all the authors.

## List of Abbreviations

ACD: Advanced Cell Diagnostics
AD: Alzheimer’s disease
ADRC: Alzheimer’s Disease Research Center
ADRD: Alzheimer’s disease and related dementias
BBDP: Brain Bank Donation Program
BSA: Bovine serum-albumin
CCK: Cholecystokinin
CGM: Central gray of the medulla
CRH: Corticotrophin releasing hormone
CRHR1: Corticotrophin releasing hormone receptor 1
DAB: 3,3′-Diaminobenzidine
D2R: D2-like dopamine receptor
EP: Ependymal
GABA: γ-Aminobutyric acid
GAD: Glutamic acid decarboxylase
H&E: Hematoxylin and Eosin
HG: Hypocellular gap
IHC: Immunohistochemistry
IR: Immunoreactivity
ISH: *In situ* hybridization
LC: Locus coeruleus
MAP2: Microtubule-associated protein-2
MCH: Melanin-concentrating hormone
MCH-1: Melanin-concentrating hormone receptor-1
mlf: Medial longitudinal fasciculus
NMB: Neuromedin-B
NI: Nucleus incertus
ON: Overnight
OX1: Orexin receptor-1
OX2: Orexin receptor-2
PAG: Periaqueductal gray
PBS: Phosphate buffered saline
PBS-T: Phosphate-buffered saline with Tween^®^ 20 detergent
PRn: Pontine raphe nucleus
RLN3: Relaxin-3
RT: Room temperature
RXFP3: Relaxin-family peptide-3 receptor
SHS: Septohippocampal system
SN: Substantia nigra
TBS-T: Tris-buffered saline with 0.1% Tween^®^ 20 detergent
vGAT1: Vesicular GABA transporter-1
4V: Fourth ventricle
5-HT1A: Serotonin-1A receptor

## Acknowledgments

The authors declare no competing financial interests. The authors are grateful to the Banner Sun Health Research Institute Brain and Body Donation Program of Sun City, Arizona for the provision of human biological materials.

## Figure Legends

**Supplementary Fig. 1.**
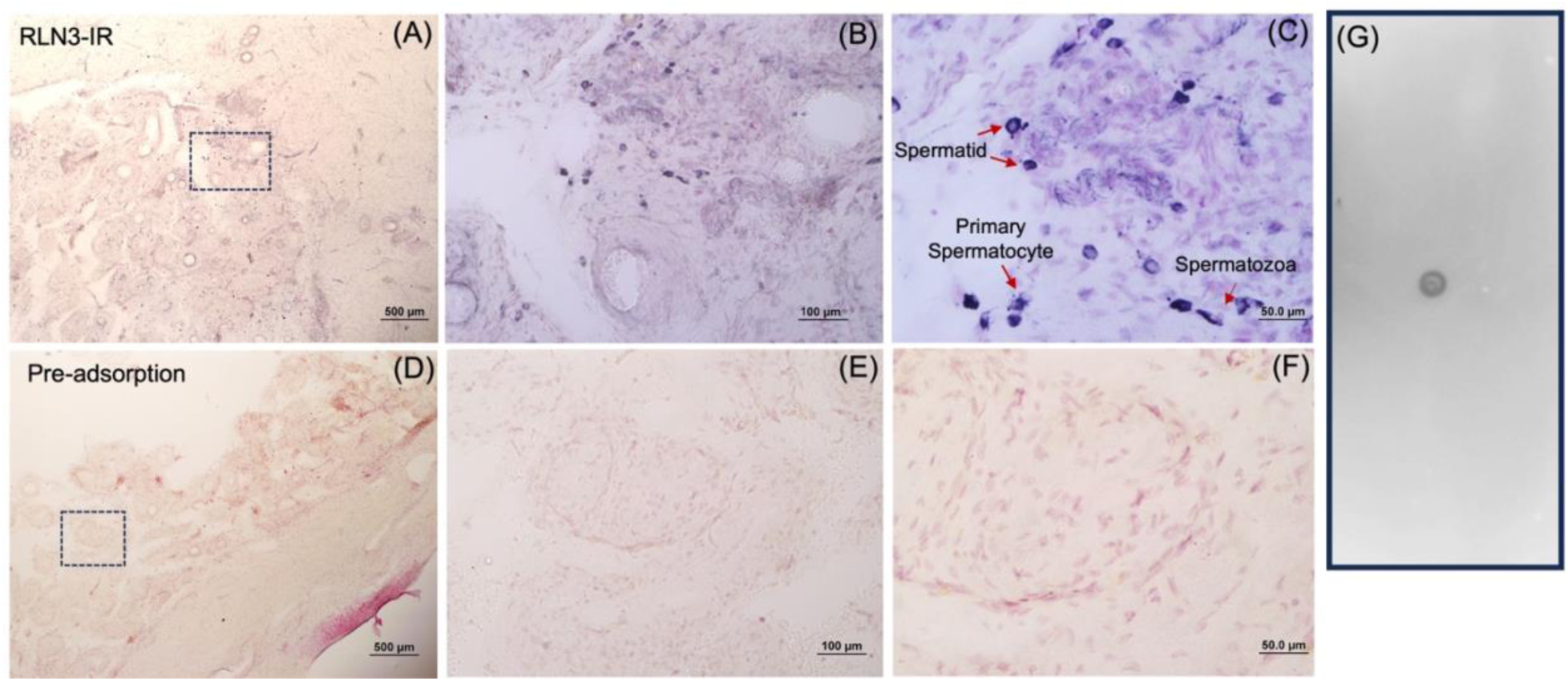
Specificity of RLN3 Antiserum Assessed in Human Testis. (**A-C)** RLN3 immunoreactivity (IR) detected in spermatozoa cells in the human testis; (**B,C)** Higher magnification images of RLN3-IR in boxed area in A. (**D-F)** Pre-adsorption of RLN3 antibody pre-incubated overnight with RLN3 recombinant peptide. (**E,F)** Higher magnification images of boxed area in D. Tissue counterstained with neutral red. (**G)** Positive dot-blot for RLN3 antiserum against native peptide; circular immunoreactivity in the center of the cellulose membrane reflects the binding of the peptide (RLN3) to the antibody (anti-RLN3).

**Supplementary Fig. 2.**
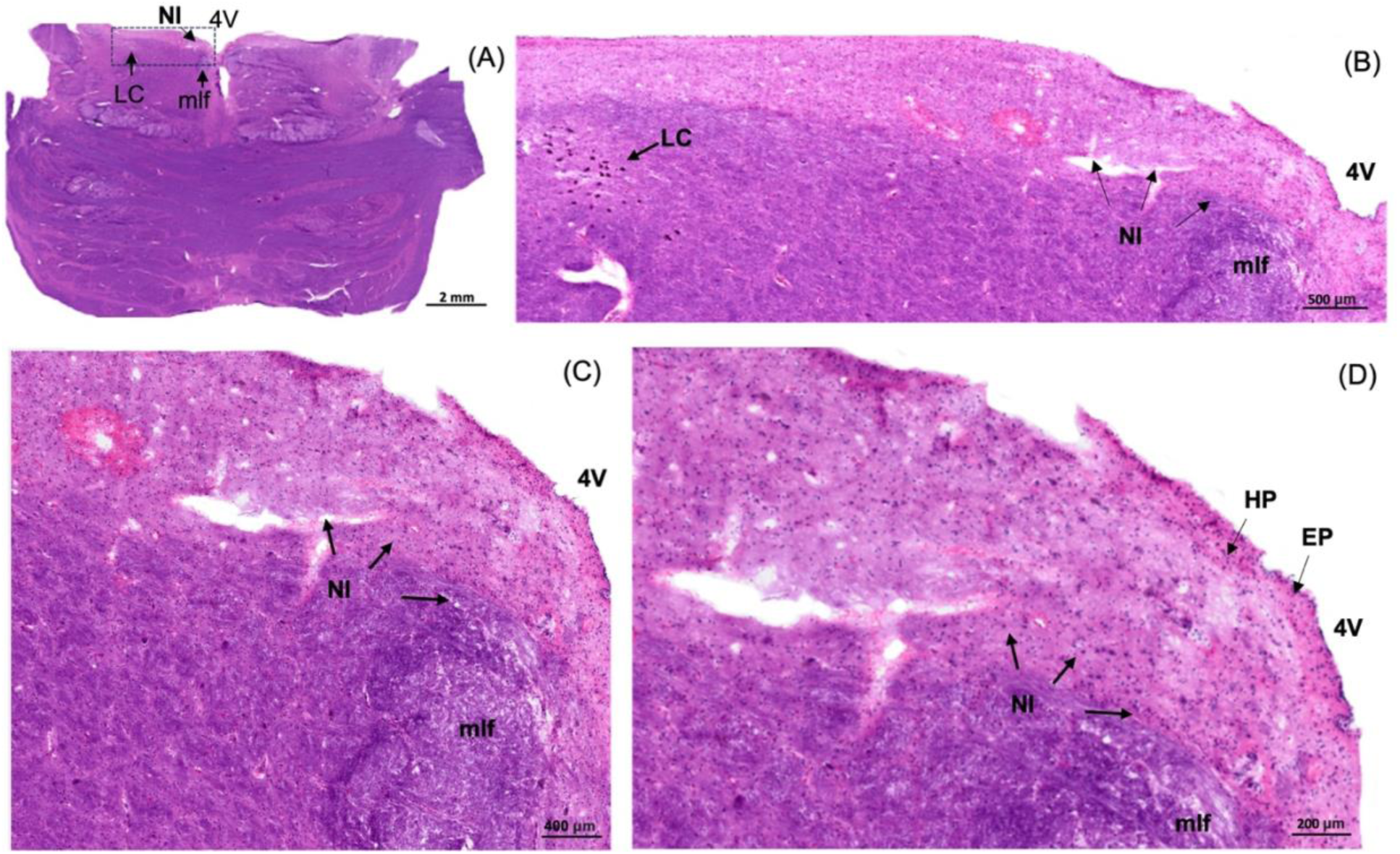
Hematoxylin and Eosin Staining of the Coronal Pons of Human Brain. (**A**) Low-power image of the neuroanatomy at the pons containing the NI (**B)** Higher magnification of the boxed area in A. (**C)** Higher magnification of the boxed area in B. (**D)** Higher magnification of the boxed area in C. Abbreviations: EP, ependyma, 4V, fourth, ventricle, HG, Hypocellular gap, LC, locus coeruleus, mlf, medial longitudinal fasciculus, NI, nucleus incertus.

